# Sphingolipid Hydroxylation Governs Fungal Pathogenicity and Drug Susceptibility: Scs7 as a Druggable Target in *Candida albicans*

**DOI:** 10.64898/2026.11.09.765499

**Authors:** Shenlong Mo, Xiaoyan Chen, Boming Yu, Jia Peng, Mengmin Zhan, Qianjun Zhao, Zhitao Deng, Lingbing Zeng, Xiaotian Huang, Jingjing Duan

## Abstract

Sphingolipids are essential structural components of the fungal plasma membrane and key signaling hubs that orchestrate pathogenicity. While both fungi and mammals perform sphingolipid hydroxylation, they rely on evolutionarily divergent enzymatic systems, establishing a biochemical divergence exploitable for selective targeting. Here, through a screening of sphingolipid metabolic genes, we identify the sphingolipid α-hydroxylase Scs7 as a central regulator of membrane homeostasis, virulence, and caspofungin susceptibility in *Candida albicans*. Genetic ablation of *SCS7* depletes hydroxylated sphingolipids, disrupts memberane organization, and compromises cell wall integrity, which collectively impair hyphal morphogenesis, biofilm formation, and stress tolerance through blockade of the cAMP-PKA signaling axis. In a murine cutaneous infection model, *SCS7* deletion markedly attenuates fungal burden, hyperinflammation, and epidermal barrier disruption, underscoring its non-redundant role in host-pathogen interaction. Leveraging the fungus-specific architecture of Scs7, we performed virtual screening and identified resveratrol as a natural Scs7 inhibitor. Resveratrol recapitulates the lipidomic and phenotypic defects of SCS7 deletion and synergizes with caspofungin to enhance fungal killing. Our study establishes sphingolipid α-hydroxylation as a master determinant of fungal membrane fitness and validates Scs7 as a promising, fungus-specific antifungal target with dual anti-virulence and chemosensitizing activities.

**Graphical Abstract:** 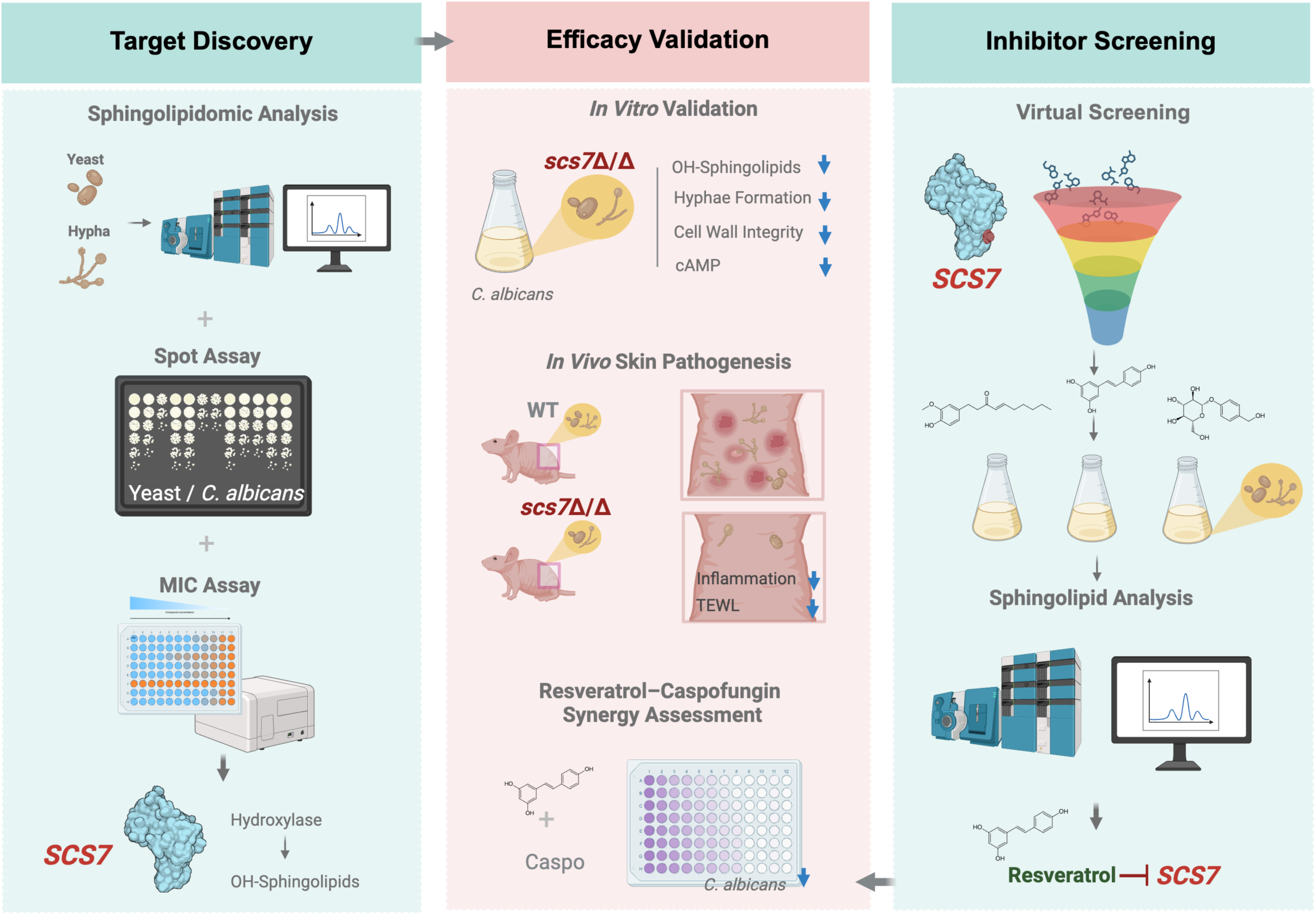

## Introduction

*Candida albicans* is classified by the World Health Organization in the critical priority group of fungal pathogens and represents one of the most prevalent commensal fungi in the human mycobiome [1–3]. In healthy individuals, this organism typically colonizes the skin, oral cavity, gastrointestinal tract, and urogenital mucosal surfaces as a low-abundance, harmless yeast, maintained in a commensal state through competition with the resident microbiota and host immune surveillance [4,5]. Upon disruption of the skin barrier, dysbiosis of the microbiota, or impairment of host immunity, *C. albicans* can rapidly transition from commensalism to pathogenicity, giving rise to a spectrum of infections characterized by erythema, erosion, pruritus, and pain. In intertriginous regions, secondary bacterial infection is common, and severe cases can progress to chronic ulceration, substantially compromising quality of life [5]. Critically, such cutaneous foci can serve as portals for bloodstream dissemination and deep-tissue invasion [6]. Beyond mucocutaneous disease, *C. albicans* also causes life-threatening invasive infections, with persistently high mortality rates in immunocompromised populations.

The current clinical antifungal armamentarium is extremely limited, consisting primarily of azoles, echinocandins, and polyenes, all of which target components of the fungal cell membrane or cell wall [7–11]. Widespread and prolonged use of these agents over the past decades has driven the emergence and global dissemination of drug-resistant *C. albicans* isolates, leading to increasingly frequent treatment failure [7,12–15]. Compounding this challenge, *C. albicans* readily forms biofilms on mucosal surfaces, which further complicates clinical management [16,17]. The biofilm extracellular matrix acts as a physical barrier that limits drug penetration and shelters large numbers of drug-tolerant persister cells, making infections difficult to eradicate, prolonging treatment duration, and further fueling the emergence and spread of resistant strains [18,19]. Therefore, dissecting the core regulatory circuits that govern drug susceptibility and identifying new targets capable of potentiating existing antifungals are essential steps toward overcoming clinical resistance and developing effective synergistic therapeutic strategies.

Fungal sphingolipids have been demonstrated to be intimately linked to drug resistance and pathogenicity, and the sphingolipid metabolic pathway has emerged as a promising antifungal target with substantial translational potential [20,21]. Unlike mammals, *C. albicans* synthesizes abundant phytoceramide and diverse hydroxylated sphingolipid species, yielding a sphingolipid profile that is distinctive of this fungus [22–25]. Targeting this fungus-enriched modification offers the possibility of selective intervention without broadly disrupting host metabolism. Notably, drug-resistant clinical isolates exhibit profoundly remodeled sphingolipid profiles; in *Candida auris* strains with stable fluconazole resistance, phytoceramide and hydroxylated sphingolipid levels are markedly elevated [26,27]. Moreover, alterations in sphingolipid composition per se can induce fluconazole resistance in *C. albicans* [28]. Although sphingolipid metabolism has been extensively explored as an antifungal target, owing to the complex regulatory networks and diverse metabolic branches of this pathway, the specific mechanisms linking sphingolipid metabolism to drug resistance and pathogenicity remain incompletely understood, leaving numerous gaps to be addressed.

In this study, we screened sphingolipid metabolism-related genes for their roles in drug susceptibility in *C. albicans* and identified *SCS7* as a key gene governing caspofungin resistance and virulence. We demonstrate that *SCS7* modulates these phenotypes by controlling cell wall integrity, biofilm formation, and the cAMP-PKA signaling pathway. Furthermore, through virtual screening, we identified candidate small molecules that can be used in combination with caspofungin to synergistically overcome drug resistance. Collectively, our findings establish *SCS7* as a central regulator in sphingolipid metabolism-mediated drug resistance and virulence in *C. albicans*, providing a theoretical basis and preclinical experimental support for developing novel antifungal strategies that target sphingolipid metabolism, with important implications for the management of drug-resistant *C. albicans* infections.

## Results

### Sphingolipid remodeling during hyphal development and identification of *SCS7* via genetic screening

Sphingolipids are core structural components of the fungal cell membrane and key signaling molecules, broadly participating in morphogenesis, stress responses, and virulence regulation. Concurrently, sphingolipid metabolic pathways have gained extensive attention as promising targets for antifungal sensitization. The yeast-to-hypha transition in *Candida albicans* is a core virulence determinant. To dissect the dynamic features of sphingolipid metabolism during this process, we performed quantitative sphingolipidomic analysis on yeast-form and Spider medium-induced hyphal *C. albicans* (Figure 1A). Results revealed an elevation of total ceramides in hyphal cells, encompassing dihydroceramides (d18:0, d20:0) and phytoceramides (t18:0, t20:0) (Figure 1B). Further analysis by fatty acid chain length demonstrated a consistent increase across various ceramide species (Figure S1A–D), suggesting global activation of the sphingolipid synthesis pathway during hyphal formation. Additionally, targeted analysis of a mouse systemic kidney infection transcriptomic dataset (GSE327269) revealed significant enrichment of sphingolipid metabolic pathways at 48 h post-infection, with multiple key synthase genes progressively upregulated (Figure S1E–F). These *in vivo* data further corroborate the close association between sphingolipid metabolic reprogramming and the pathogenic process of *C. albicans*.

**Figure 1.**
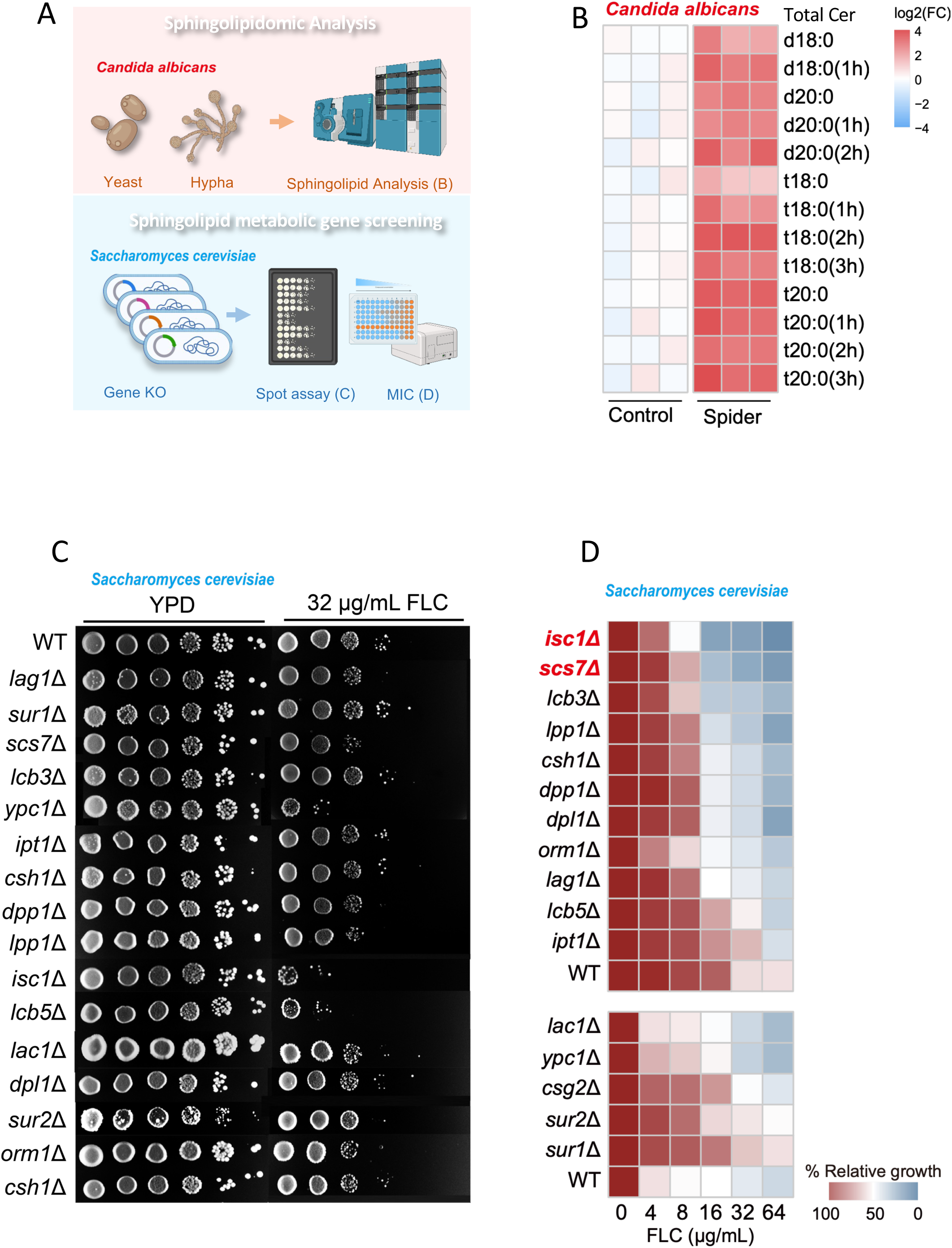
Sphingolipid metabolic reprogramming accompanies *Candida albicans* morphogenesis, and genetic screening in *Saccharomyces cerevisiae* identifies key regulators of antifungal susceptibility (A) Schematic of the experimental workflow. Sphingolipidomic analysis was performed on *C. albicans* yeast-form and hyphae cells (top). A sphingolipid metabolic gene deletion library in *S. cerevisiae* was screened via spot assay and MIC determination (bottom). (B) Heatmap showing total ceramide (Total Cer) levels in *C. albicans* yeast (Control) and hyphae (Spider). Color intensity represents log_2_ fold change (FC) (red, upregulated; blue, downregulated). (C) Spot assay comparing the growth of *S. cerevisiae* wild-type (WT) and sphingolipid metabolic gene deletion mutants on YPD with or without 32 μg/mL fluconazole (FLC). (D) Heatmap of relative growth of *S. cerevisiae* mutants across different FLC concentrations (0 to 64 μg/mL).

To further identify key functional genes regulating antifungal susceptibility in the sphingolipid metabolic pathway, we employed *Saccharomyces cerevisiae* as a model organism for screening, which has a mature genetic manipulation system and a highly conserved sphingolipid pathway. We constructed a deletion mutant library covering 16 key sphingolipid metabolic genes and employed a two-step strategy (spot dilution assay-based primary screening followed by MIC quantitative validation) to identify genes whose deletion enhances susceptibility to fluconazole (FLC) (Figure 1A). Multiple mutants exhibited increased FLC susceptibility (Figure 1C). Following further MIC quantitative validation, *scs7*Δ and *isc1*Δ predominantly displayed stable and significant FLC-sensitive phenotypes (Figure 1D), and were thus selected as core candidate genes for subsequent validation of their dual regulatory functions in virulence and drug responses in *C. albicans*.

### *SCS7* regulates caspofungin susceptibility and is required for morphogenesis and infection in *C. albicans*

To validate whether the drug susceptibility phenotypes observed in *S. cerevisiae*are conserved in *C. albicans*, we constructed homozygous deletion strains of *SCS7* and *ISC1* in *C. albicans* (*scs7*Δ/Δ, *isc1*Δ/Δ). Successful generation of the mutants was confirmed by transcriptional analysis (Figure S2A, B). We then performed systematic antifungal susceptibility testing with three common clinical antifungal drugs: caspofungin (CAS), 5-fluorocytosine (5-FC), and fluconazole (FLC). The *scs*7Δ/Δ strain exhibited significantly increased susceptibility to CAS but showed no significant changes to FLC or 5-FC. In contrast, *isc1*Δ/Δ displayed no stable susceptibility changes to any of the three drugs (Figure 2A). Furthermore, complementation of *SCS7* (*scs7*Δ/*SCS7*) restored CAS susceptibility to wild-type levels (Figure 2B, Figure S2C), confirming *SCS7* as a key regulator of CAS susceptibility in *C. albicans*.

**Figure 2.**
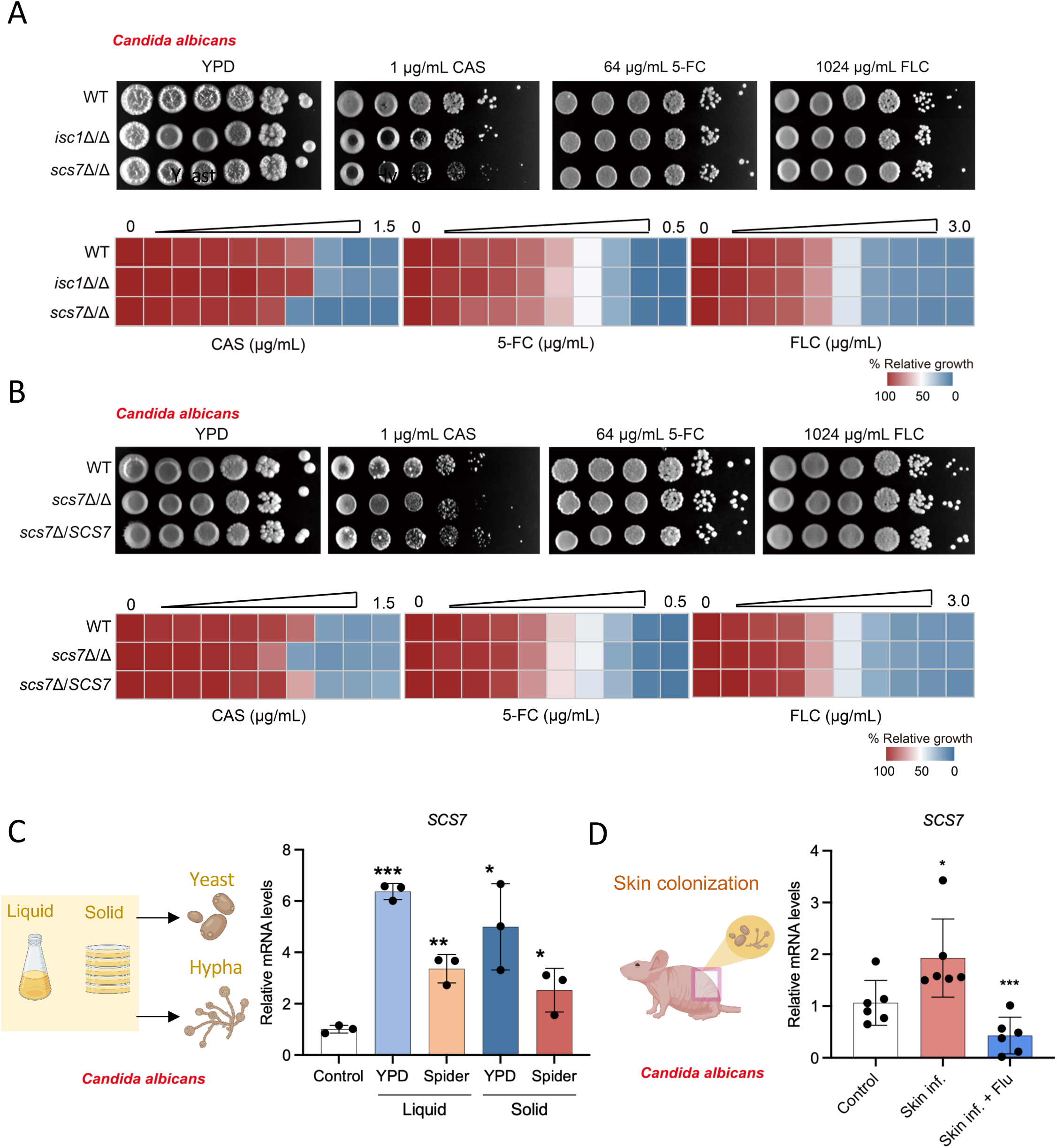
*SCS7* regulates caspofungin susceptibility and is associated with morphogenesis and infection in *C. albicans*. (A) Susceptibility analysis of *C. albicans scs7*Δ/Δ and *isc1*Δ/Δ deletion mutants to caspofungin (CAS), 5-fluorocytosine (5-FC), and fluconazole (FLC). Spot assay (top) and relative growth heatmap (bottom) comparing WT, *scs7*Δ/Δ, and *isc1*Δ/Δ strains under CAS (1 μg/mL), 5-fluorocytosine (5-FC, 64 μg/mL), or fluconazole (FLC, 1024 μg/mL) stress. (B) Complementation analysis. Same experimental design as in (A), comparing WT, *scs7*Δ/Δ, and *scs7*Δ/*SCS7* strains. (C) Relative mRNA expression levels of *SCS7* in *C. albicans* yeast (liquid and solid YPD) and hyphae (liquid and solid Spider medium), normalized to 18S rRNA. (D) *In vivo* infection model. Six-to eight-week-old female BALB/c-nu nude mice (n = 6) underwent tape-stripping on the back skin on day 0, followed by topical inoculation of fungi on days 0, 2, 4, and 6. Dorsal skin tissues were collected on day 14 for subsequent analyses. Relative mRNA expression levels of *SCS7* in *C. albicans* from uninfected mouse skin (Control) and infected mouse skin on day 14 (Skin inf.), with or without fluconazole treatment (Skin inf. + Flu), normalized to 18S rRNA. Data are presented as mean ± SD. Statistical analyses were performed using two-tailed unpaired t-tests for (C) and (D). Significance levels are defined as \**p* < 0.05, \*\**p* < 0.01, and \*\*\**p* < 0.001, with a sample size of n = 3 for (C) and n = 6 for (D).

On the other hand, transcriptional regulation analysis revealed that *SCS7* expression is tightly associated with the yeast-to-hypha transition and infection progression in *C. albicans*. In both liquid and solid Spider media, *SCS7* mRNA levels in hyphal cells were significantly higher than in yeast cells (Figure 2C), highly consistent with the activated sphingolipid synthesis observed in the sphingolipidomic analysis. In a mouse skin colonization infection model, *SCS7* expression was significantly upregulated in *C. albicans* at the infected sites (Figure 2D). Collectively, these results demonstrate that *SCS7* not only specifically regulates caspofungin susceptibility in *C. albicans* but also plays a critical role in hyphal morphogenesis and in vivo infection progression, suggesting its dual involvement in both fungal virulence and antifungal drug responses.

### *SCS7* deletion reduces hydroxylated sphingolipid levels in *C. albicans*

*SCS7* encodes a fatty acid α-hydroxylase that catalyzes hydroxylation at the sphingolipid fatty acid chain (Figure 3A) [22]. To determine the impact of *SCS7* deletion on hydroxylated sphingolipid levels in *C. albicans*, we performed quantitative sphingolipidomic profiling of wild-type (WT), *scs7*Δ/Δ mutant, and *scs7*Δ/*SCS7* complemented strains by liquid chromatography–mass spectrometry. Compared with WT, the *scs7*Δ/Δ mutant displayed a pronounced depletion of ceramide species bearing hydroxylated fatty acid chains (including d18:0-Cer, d20:0-Cer, t18:0-Cer, and t20:0-Cer), concomitant with a corresponding increase in their non-hydroxylated counterparts.

**Figure 3.**
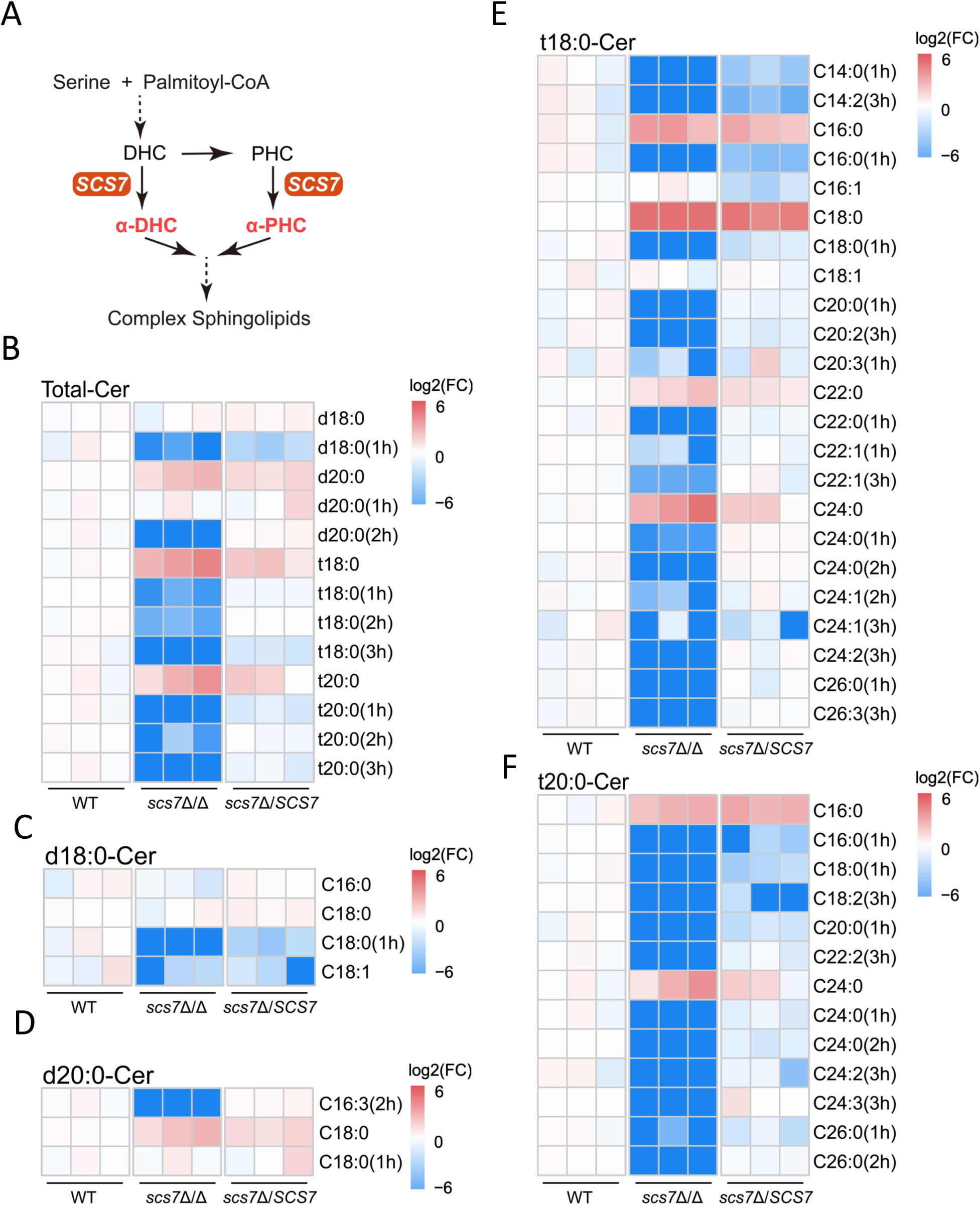
***SCS7* deletion reduces levels of hydroxy ceramides in *C. albicans*.** (A) Schematic of the sphingolipid metabolic pathway in *C. albicans*. The enzyme encoded by *SCS7* catalyzes α-hydroxylation of the fatty acid chain of ceramides, generating α-OH dihydroceramide (α-DHC) and α-OH phytoceramide (α-PHC). (B) Heatmap showing the abundance of ceramide species in wild-type (WT), *scs7*Δ/Δ deletion mutant, and *scs7*Δ/*SCS7* complemented strains. Ceramides are classified by long-chain base type, dihydrosphingosine (d-18:0 and d-20:0) or phytosphingosine (t-18:0 and t-20:0), and fatty acid hydroxylation status (1h/2h/3h, indicating the number of hydroxyl groups). (C-F) Heatmaps showing the abundance of species with different acyl chain lengths in wild-type (WT), *scs7*Δ/Δ deletion mutant, and *scs7*Δ/*SCS7* complemented strain. (C) d18:0 dihydroceramide. (D) d20:0 dihydroceramide. (E) t18:0 phytoceramide. (F) t20:0 phytoceramide. 1h, 2h, and 3h denote the number of hydroxyl groups on the fatty acid chain. All heatmap data are presented as log₂ fold change (FC) relative to wild-type (WT) control, with red indicating increased abundance and blue indicating decreased abundance compared to WT.

In the *scs7*Δ/*SCS7* complemented strain, the levels of these hydroxylated ceramides were largely restored to WT levels (Figure 3B). Further analysis across different fatty acid chain lengths confirmed a pervasive reduction in hydroxylated sphingolipid species in the *scs7*Δ/Δ mutant (Figures 3B–F). Collectively, these data demonstrate that loss of *SCS7* in *C. albicans* triggers a remodeling of the sphingolipid profile, underscoring a potential role for hydroxylated sphingolipids in mediating drug resistance and host infection.

### *SCS7* deletion disrupts cell wall integrity and attenuates virulence

The fungal cell wall is a critical barrier against environmental insults, and sphingolipid metabolism underpins its homeostasis [16,17]. To assess whether *SCS7* is required for cell wall integrity in *C. albicans*, we examined sensitivity to Congo red (CR) and sodium dodecyl sulfate (SDS). CR functions as a cell wall-perturbing agent that compromises growth in a PKC (MAPK) pathway-dependent manner, whereas SDS directly disrupts cell wall and membrane architecture. Under non-stress conditions, WT, *scs7*Δ/Δ, and *scs7*Δ/*SCS7* strains grew comparably. However, in the presence of CR or SDS, growth of the *scs7*Δ/Δ mutant was profoundly impaired, and this inhibition was partially rescued in the complemented strain (Figure 4A), indicating that loss of *SCS7* compromises cell wall integrity. We next asked whether cell wall composition was altered. Calcofluor white (CFW) staining revealed comparable chitin levels between the *scs7*Δ/Δ mutant and the WT strain (Figure 4B), indicating that *SCS7* deficiency primarily disrupts the overall structural integrity of the cell wall architecture rather than chitin biosynthesis.

**Figure 4.**
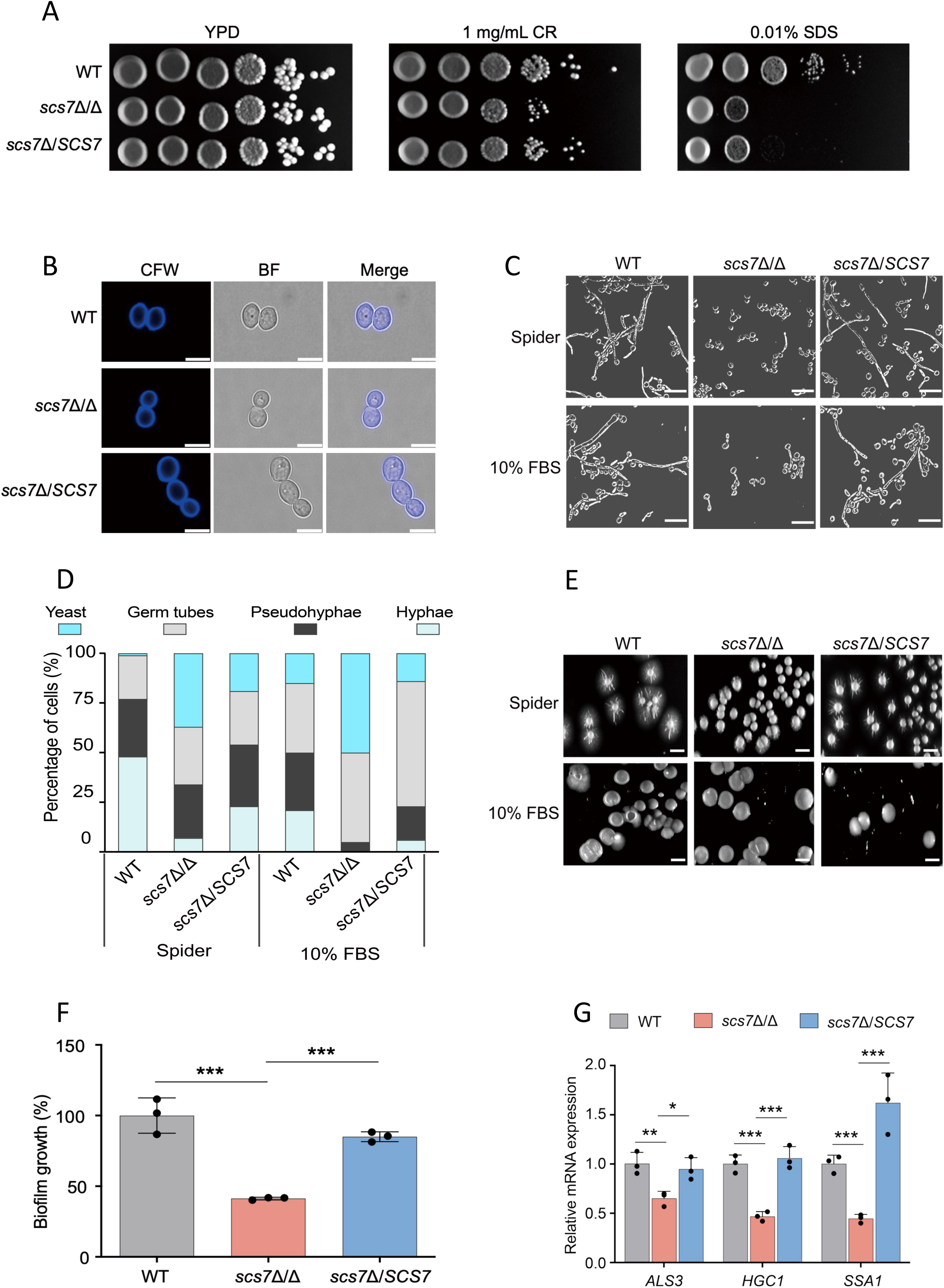
*SCS7* deletion impairs cell wall integrity, hyphal formation, and biofilm formation in *C. albicans*. (A) Spot assay detecting growth differences of *C. albicans* WT, *scs7*Δ/Δ, and *scs7*Δ/*SCS7* under cell wall stress conditions (1 mg/mL Congo red or 0.01% w/v SDS). (B) Fluorescence microscopy images after CFW staining showing chitin distribution in the cell walls of *C. albicans* WT, *scs7*Δ/Δ, and *scs7*Δ/*SCS7*; scale bar = 5 μm. (C-E) Hyphal formation of *C. albicans* induced at 37°C in RPMI 1640 with 10% FBS and in Spider liquid medium for 24 h (C), and on solid medium for 7 days (E). (D) Classification criteria: hyphae (narrow filaments, < 2 μm wide, with parallel sides); pseudohyphae (width > 2 μm, constrictions, septum at the mother-daughter cell junction, non-parallel sides); short germ tubes (swollen, less polarized structure). At least five fields and 100 cells were classified per strain; scale bar = 10 μm (C); 1 cm (E). (F) Biofilm formation of *C. albicans* WT, *scs7*Δ/Δ, and *scs7*Δ/*SCS7* after 24 h at 37°C, compared by crystal violet staining. (G) Relative mRNA expression levels of hyphal development and virulence-related genes ALS3, HGC1, and SSA1 in *C. albicans* WT, *scs7*Δ/Δ, and *scs7*Δ/*SCS7* strains, normalized to 18S rRNA. Data are presented as mean ± SD and were analyzed by one-way ANOVA followed by Dunnett’s multiple comparisons test (F) and two-tailed unpaired *t*-tests (G). Significance levels: \**p* < 0.05, \*\**p* < 0.01, \*\*\**p* < 0.001, with a sample size of n = 3.

The yeast-to-hypha transition is a prerequisite for biofilm formation and host invasion in *C. albicans* [18,29]. We therefore evaluated the impact of *SCS7* on morphogenesis by inducing hyphae in Spider medium and in RPMI 1640 supplemented with 10% FBS, and by monitoring colony morphology on solid inducing media. In both liquid Spider medium and RPMI 1640 supplemented with 10% FBS, the *scs7*Δ/Δ mutant exhibited a pronounced defect in hyphal development, whereas the complemented strain partially restored hypha formation (Figures 4C, D). On solid Spider agar, colonies of the *scs7*Δ/Δ mutant displayed smooth borders and markedly impaired filamentous extension (Figure 4E). Consistent with the hyphal defect, the *scs7*Δ/Δ mutant displayed a marked deficit in adhesion to abiotic surfaces and diminished biofilm biomass (Figure 4F), accompanied by downregulation of adhesion- and hypha-associated genes such as *ALS3* and *HGC1* (Figure 4G). Collectively, these data demonstrate that *SCS7* deletion severely compromises cell wall integrity and impairs key virulence attributes in *C. albicans*, including hyphal morphogenesis, substrate adhesion, and biofilm formation.

### *SCS7* deletion attenuates the skin infection capacity of *Candida albicans*

To define the role of *SCS7* in *C. albicans* cutaneous infection, we employed a BALB/c nude mouse model of cutaneous candidiasis. Wild-type (WT), *scs7*Δ/Δ mutant, and *scs7*Δ/*SCS7* complemented strains were inoculated onto the dorsal skin, and disease progression was monitored over 14 days (Figure 5A). Mice infected with WT or complemented strains developed characteristic signs of cutaneous candidiasis, including pronounced erythema, edema, and desquamation. In stark contrast, infection with the *scs7*Δ/Δ mutant elicited markedly attenuated skin lesions (Figure 5B), indicating that *SCS7* deletion compromises the cutaneous virulence of *C. albicans*.

**Figure 5.**
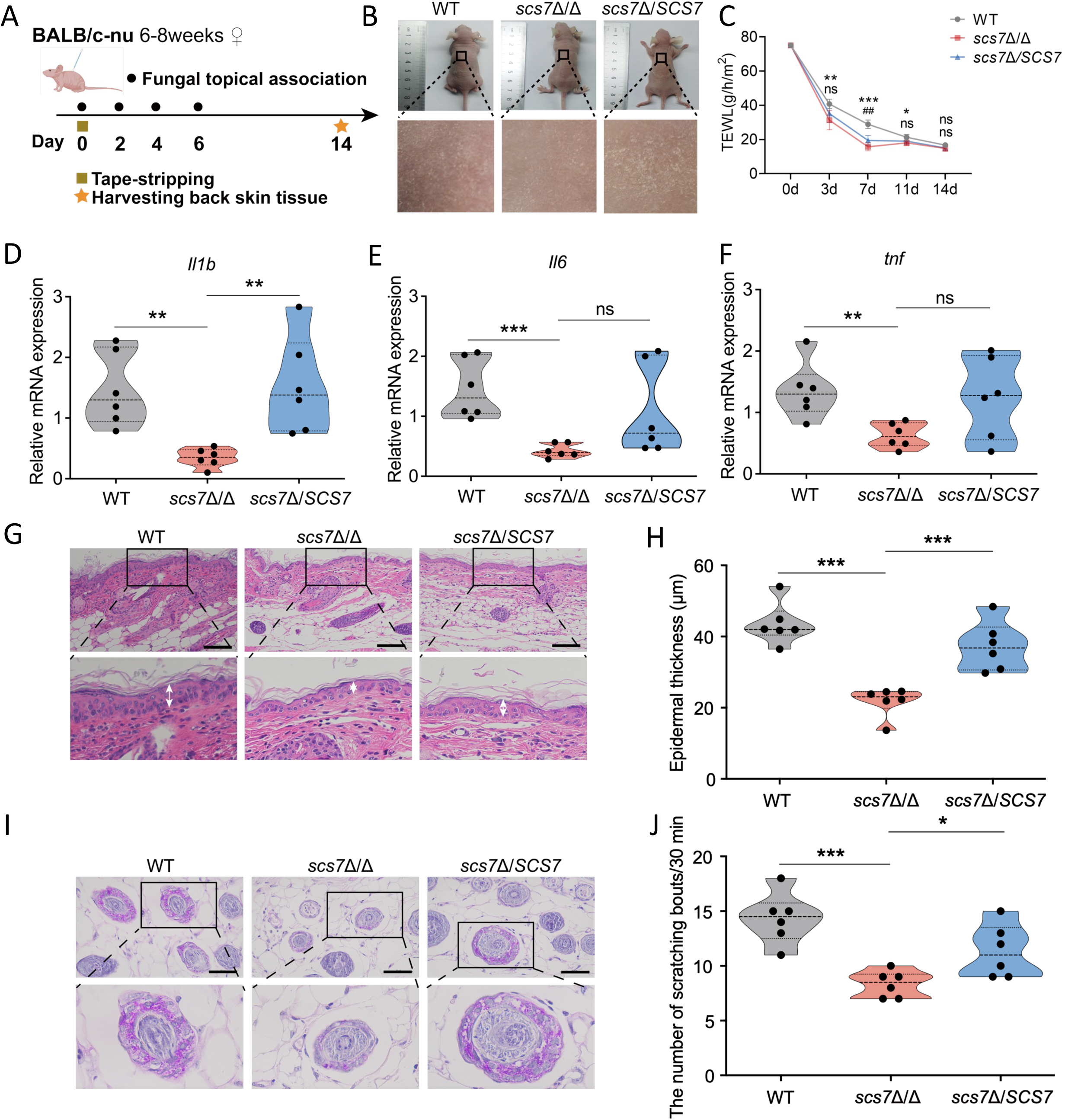
***SCS7* deletion attenuates the skin infection capacity of *C. albicans.*** (A) Schematic of the mouse skin colonization model for *C. albicans*, using six-to eight-week-old female BALB/c-nu nude mice (n = 6) Tape-stripping was performed on the back skin on day 0, followed by topical inoculation of fungi on days 0, 2, 4, and 6. Dorsal skin tissues were collected on day 14 for subsequent analyses. (B) Representative images of mouse back skin on day 14. (C) Transepidermal water loss (TEWL) measured on days 3, 7, 11, and 14. (D–F) Relative mRNA expression levels of *Il1b* (D), *Il6* (E), and *Tnf* (F) in the dorsal skin tissues of mice on day 14, determined by qPCR and normalized to *Actb*. (G–H) Histopathological examination of skin on day 14 by H&E staining (G, scale bar = 100 μm), with epidermal thickness measured at five points per mouse (H). (I) PAS staining of dorsal skin tissue sections on day 14, showing positive reactions (red) for fungal hyphae/spores in sebaceous glands; scale bar = 50 μm. (J) Behavioral recording and analysis of scratching of the back skin on day 14. A single continuous scratching bout was counted as one episode, and the total number of episodes within 30 min was recorded. Data are presented as mean ± SD. In (C), two-way ANOVA followed by Dunnett’s multiple comparisons test was used, with * indicating WT vs. *scs7*Δ/Δ and # indicating *scs7*Δ/*SCS7* vs. *scs7*Δ/Δ. For (D–F, H, J), two-tailed unpaired *t*-tests were performed. Significance is denoted as \**p* < 0.05, \*\**p* < 0.01, \*\*\**p* < 0.001, with a sample size of n = 6.

To quantitatively assess skin barrier disruption, we measured transepidermal water loss (TEWL), a well-established functional indicator of stratum corneum integrity with elevated levels signifying barrier impairment[30]. On day 7 post-infection, the WT-infected group exhibited significantly higher TEWL values than the *scs7*Δ/Δ-infected group, suggesting more severe early disruption of the skin water-retention barrier. Despite the restoration of TEWL to normal levels by day 14 (Figure 5C), skin homogenates from the WT-infected group continued to exhibit significantly higher levels of *Il1b*, *Il6* and *Tnf* than those from the *scs7*Δ/Δ group (Figures 5D–F), confirming that *SCS7* deficiency markedly attenuates the capacity of *C. albicans* to induce a pro-inflammatory response in the skin.

Histopathological analysis of infected skin further corroborated these findings. Hematoxylin and eosin (H&E) staining revealed prominent epidermal hyperplasia in WT-infected skin, whereas epidermal thickening was significantly milder in the *scs7*Δ/Δ-infected group (Figures 5G–H). Periodic acid–Schiff (PAS) staining confirmed a visibly reduced fungal burden in the skin of *scs7*Δ/Δ-infected mice relative to WT controls, indicative of impaired cutaneous colonization upon *SCS7* loss (Figure 5I). Given that pruritus correlates with infection severity, we assessed spontaneous scratching behavior on day 14. Despite the ongoing recovery of the skin barrier, scratching frequency remained significantly elevated in the WT-infected group compared to the *scs7*Δ/Δ-infected mice. This phenotype was rescued in the complemented strain, which restored scratching activity to near-WT levels (Figure 5J). Collectively, these results demonstrate that *SCS7* deletion broadly compromises the capacity of *C. albicans* to induce inflammation, disrupt the epidermal barrier, and colonize the skin during cutaneous infection, thereby markedly reducing its skin pathogenic potential.

### *SCS7* deletion suppresses hypha formation by blocking the cAMP-PKA signaling pathway

Based on the severe hyphal developmental defects observed in the *scs7*Δ/Δ mutant (Figures 4C–G), we hypothesized that the primary phenotypic consequence of *SCS7* loss is impaired hypha formation, which in turn compromises skin adhesion and colonization by *C. albicans*. To elucidate the molecular basis of this phenotype, we examined the activity of the cAMP-PKA signaling pathway, the central regulatory hub governing hyphal development in this fungus. qRT-PCR analysis revealed that, relative to the wild-type strain, the *scs7*Δ/Δ mutant displayed markedly downregulated transcription of *TPK2*, which encodes a *PKA* catalytic subunit, while *TPK1* showed a decreasing trend. The expression of the downstream transcription factors *EFG1* and *FLO8* was also significantly reduced. In the complemented strain *scs7*Δ/*SCS7*, the transcript levels of *FLO8* and *TPK2* were substantially restored (Figure 6A). Quantification of intracellular cAMP revealed a significant reduction in cAMP levels in the *scs7*Δ/Δ mutant compared to the WT strain, and levels were restored to near-WT levels in the complemented strain (Figure 6B). To determine whether cAMP deficiency directly causes the hyphal defect, we demonstrated that supplementing with exogenous cAMP rescued hyphal development in the *scs7*Δ/Δ mutant in a dose-dependent manner (Figures 6C,D). Collectively, these data demonstrate that *SCS7* deletion lowers intracellular cAMP levels, thereby blocking activation of the cAMP-PKA signaling cascade and suppressing hypha formation in *C. albicans*.

**Figure 6.**
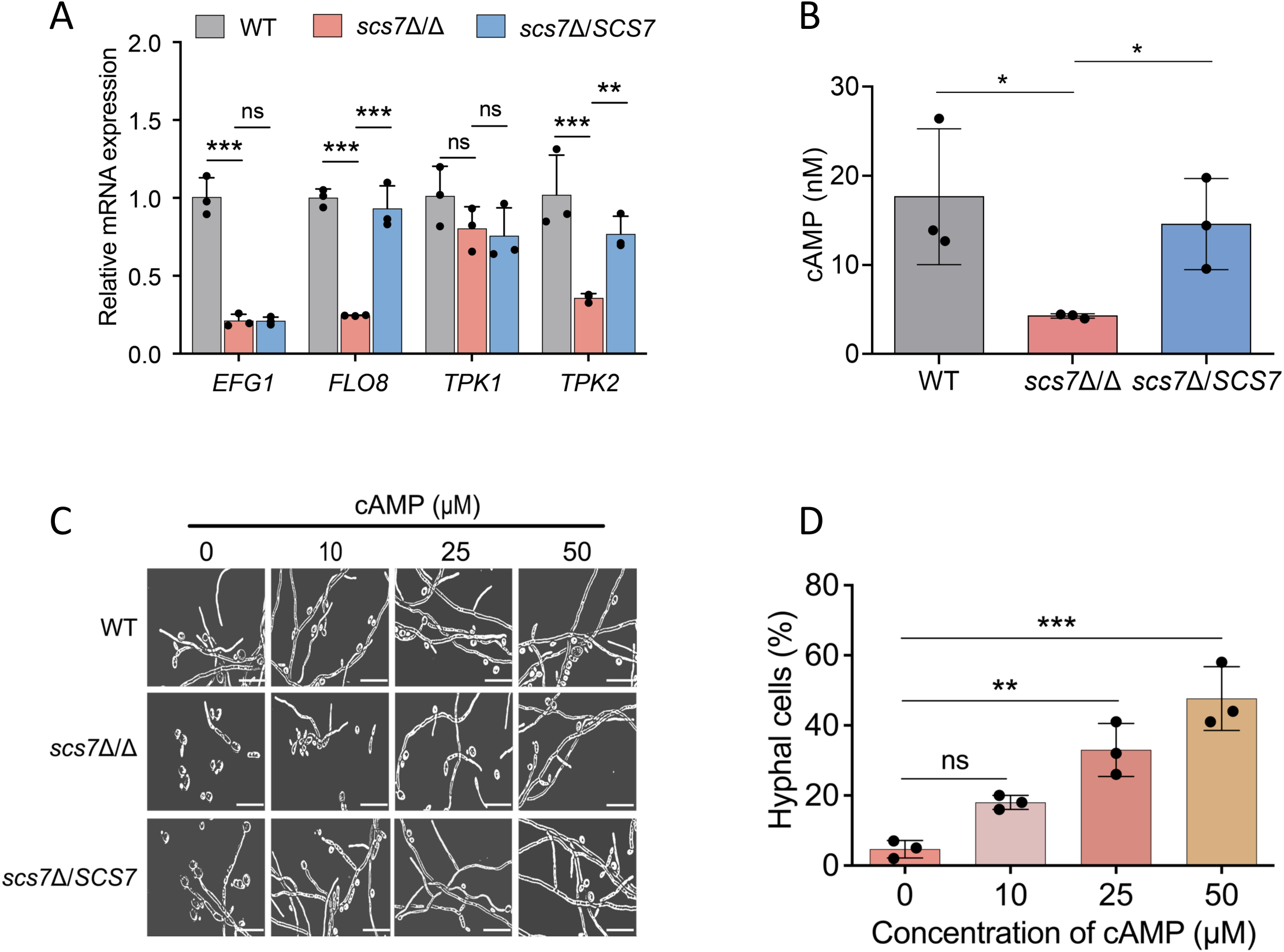
*SCS7* deletion suppresses hypha formation by blocking the cAMP-PKA signaling pathway. (A) Relative mRNA expression levels of cAMP-PKA signaling pathway-related genes (*EFG1*, *FLO8*, *TPK1*, *TPK2*) in wild-type (WT), *scs7*Δ/Δ, and *scs7*Δ/*SCS7* strains. (B) Intracellular cAMP concentrations in WT, *scs7*Δ/Δ, and *scs7*Δ/*SCS7* strains, quantified using the cAMP-Glo™ Assay kit. (C-D) Hyphal formation in WT, *scs7*Δ/Δ, and *scs7*Δ/*SCS7* strains following treatment with exogenous cAMP (0, 10, 25, or 50 μM) in Spider medium at 37°C for 24 h. Representative microscopy images are shown in (C), scale bar = 10 μm. (D) The percentage of hyphal cells was calculated by counting at least 100 cells per group.Data are presented as mean ± SD. Statistical analyses were performed using two-way ANOVA followed by Dunnett’s multiple comparisons test for (A), two-tailed unpaired t-test for (B), and one-way ANOVA followed by Dunnett’s multiple comparisons test for (D). Significance levels are defined as \**p* < 0.05, \*\**p* < 0.01, and \*\*\**p* < 0.001, with a sample size of n = 3.

### Virtual screening of natural antifungal compounds targeting Scs7

To identify natural antifungal compounds that target Scs7, we leveraged AlphaFold 3 to predict the three-dimensional (3D) structure of *C. albicans* Scs7 (Figure 7A). The pLDDT scores indicated that the residues in functional regions exceeded 90, and Ramachandran plot analysis showed that 90.9% of residues were in favored regions and 9.1% in allowed regions (Figure 7B). Using this 3D model and Maestro, we characterized the amino acid composition and spatial configuration of the catalytic active pocket (Figure 7C), which served as the ligand binding pocket for virtual screening.

**Figure 7.**
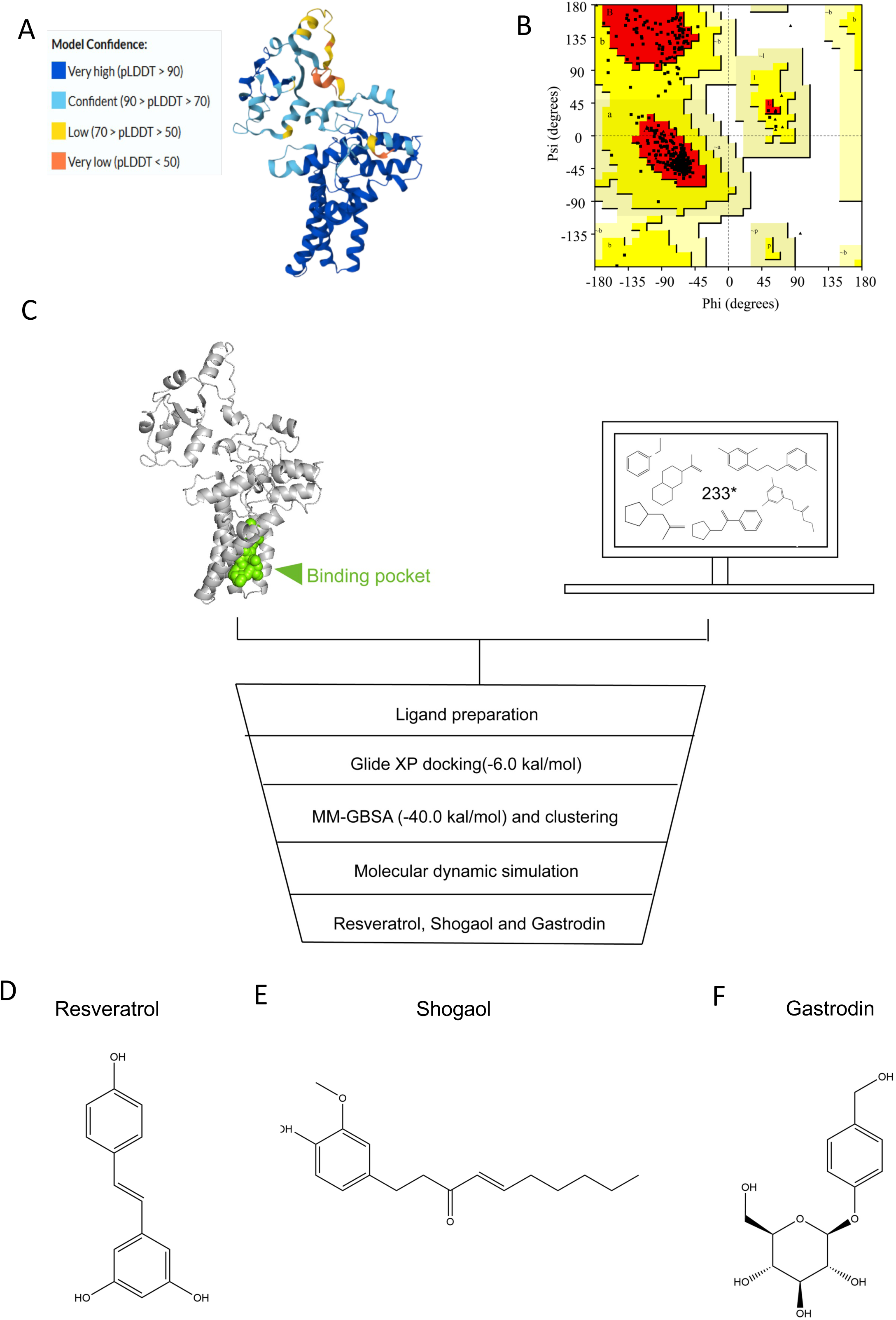
**Molecular modeling of *C. albicans* Scs7 protein and virtual screening workflow for predicting inhibitors.** (A) Predicted three-dimensional structure of *C. albicans* Scs7 generated by AlphaFold 3, colored by per-residue confidence scores pLDDT. The color scale indicates confidence levels blue for very high confidence (> 90), light blue for confident (70– 90), yellow for low confidence (50–70), and orange for very low confidence (< 50). (B) Ramachandran plot validating the stereochemical quality of the predicted Scs7 structure, showing the distribution of backbone φ/ψ dihedral angles. (C) Schematic workflow of the multi-step virtual screening strategy targeting the Scs7 binding pocket. A library of 233 antifungal compounds underwent sequential Glide XP docking, MM-GBSA rescoring and clustering and molecular dynamics simulations to assess binding stability. (D-F) Two-dimensional ligand interaction diagrams illustrating the binding modes of the three candidate inhibitors within the Scs7 binding pocket. Resveratrol (D), Shogaol (E), and Gastrodin (F) are shown with key noncovalent interactions with surrounding residues. * The 233 antifungal compounds were curated from previously published literature (see Table S5 for details).

We next applied a tiered funnel screening strategy. A total of 233 small molecules with reported antifungal activity were collected from the literature (see Table S5 for details) and subjected to structural preprocessing. All compounds were docked against the prepared Scs7 protein, and those with docking scores below −6.0 kcal/mol were retained. Candidates were further evaluated by MM-GBSA binding free energy (ΔGbind) calculations, and compounds with ΔGbind above −40 kcal/mol were eliminated. Based on docking scores, binding free energies, and binding modes, three lead compounds were selected, namely Resveratrol, Shogaol, and Gastrodin (Figure 7D).

To assess the binding stability of these three candidates, we embedded Scs7 into a POPC/DPPC lipid bilayer to mimic its native endoplasmic reticulum membrane environment and performed 300-ns molecular dynamics simulations for each complex. Analysis of Cα atom root-mean-square deviation (RMSD) showed that the Scs7-Shogaol complex remained stable at 1.0–2.0 Å during the first 250 ns, with fluctuations appearing only in the final 30 ns. The Scs7-Gastrodin complex converged rapidly at the beginning, underwent a transient fluctuation between 150 and 200 ns, and then settled at approximately 2.0 Å. Notably, although the Scs7-Resveratrol complex exhibited some fluctuations within the first 150 ns, it subsequently stabilized at 2.0–3.0 Å, with a final RMSD comparable to those of the other two complexes, indicating that Resveratrol achieves a stable binding state after an initial conformational adjustment (Figure S3A). Overall, all three candidates formed stable complexes with Scs7. Root-mean-square fluctuation (RMSF) analysis indicated that the core structures of all three complexes were fairly stable, with pronounced fluctuations only at the N- and C-terminal residues (Figure S3B).

Radius of gyration (Rg) analysis further demonstrated that the Scs7-Shogaol complex maintained a consistently low Rg, indicative of a compact, native-like fold. The Rg of the Resveratrol complex increased slightly in the early phase but gradually returned and stabilized, reflecting a reversible, localized conformational relaxation upon ligand engagement (Figure S3C). Solvent-accessible surface area (SASA) analysis showed that the SASA of the Resveratrol complex decreased continuously in the later stage and converged with that of the Shogaol system, indicating progressive burial of core residues (Figure S3D). Collectively, these metrics reveal that, despite a transient conformational adjustment, Resveratrol ultimately forms a stable and compact complex with Scs7.

Finally, we elucidated the atomic details of protein-ligand recognition. Resveratrol forms hydrogen bonds with Leu226 and Tyr317 of Scs7, with Phe293 mediating dominant hydrophobic contacts that are further stabilized through cooperative interactions with Ile297, Phe301, Phe310, Ile233, and Met229, along with water-bridged interactions. This integration of a robust hydrogen-bond network and hydrophobic packing likely confers a unique binding mode. Shogaol contributes few hydrogen bonds (only Gly313/Phe310) but establishes a strong hydrophobic core through Trp234 and Ile297. Gastrodin forms high-frequency hydrogen bonds with Asn207 and Thr314, engages a water bridge via Tyr294, and is supported by extensive hydrophobic contacts (Figures S3E–J). These results demonstrate that all three candidates achieve stable binding within the catalytic active center of Scs7 through integrated hydrogen-bond networks and hydrophobic interactions.

### Resveratrol as a promising antifungal candidate targeting Scs7 against *Candida albicans*

To evaluate the on-target inhibitory effects of the three candidate compounds on Scs7, we exposed *C. albicans* to 30 μg/mL Resveratrol, Shogaol, or Gastrodin and subsequently quantified dihydroceramide and phytoceramide species by LC-MS/MS. Compared with Shogaol and Gastrodin, Resveratrol induced a profound global depletion of sphingolipid levels (Figures 8A–E), with hydroxylated ceramides displaying the most pronounced decrease. Notably, the changes in non-hydroxylated sphingolipid species were modest. This lipid remodeling signature, characterized by a selective loss of hydroxylated sphingolipids, closely mirrored the sphingolipid profile of the *scs7*Δ/Δ mutant. These results strongly suggest that Resveratrol exerts its effects via inhibition of Scs7 to reduce hydroxylated sphingolipid synthesis.

**Figure 8.**
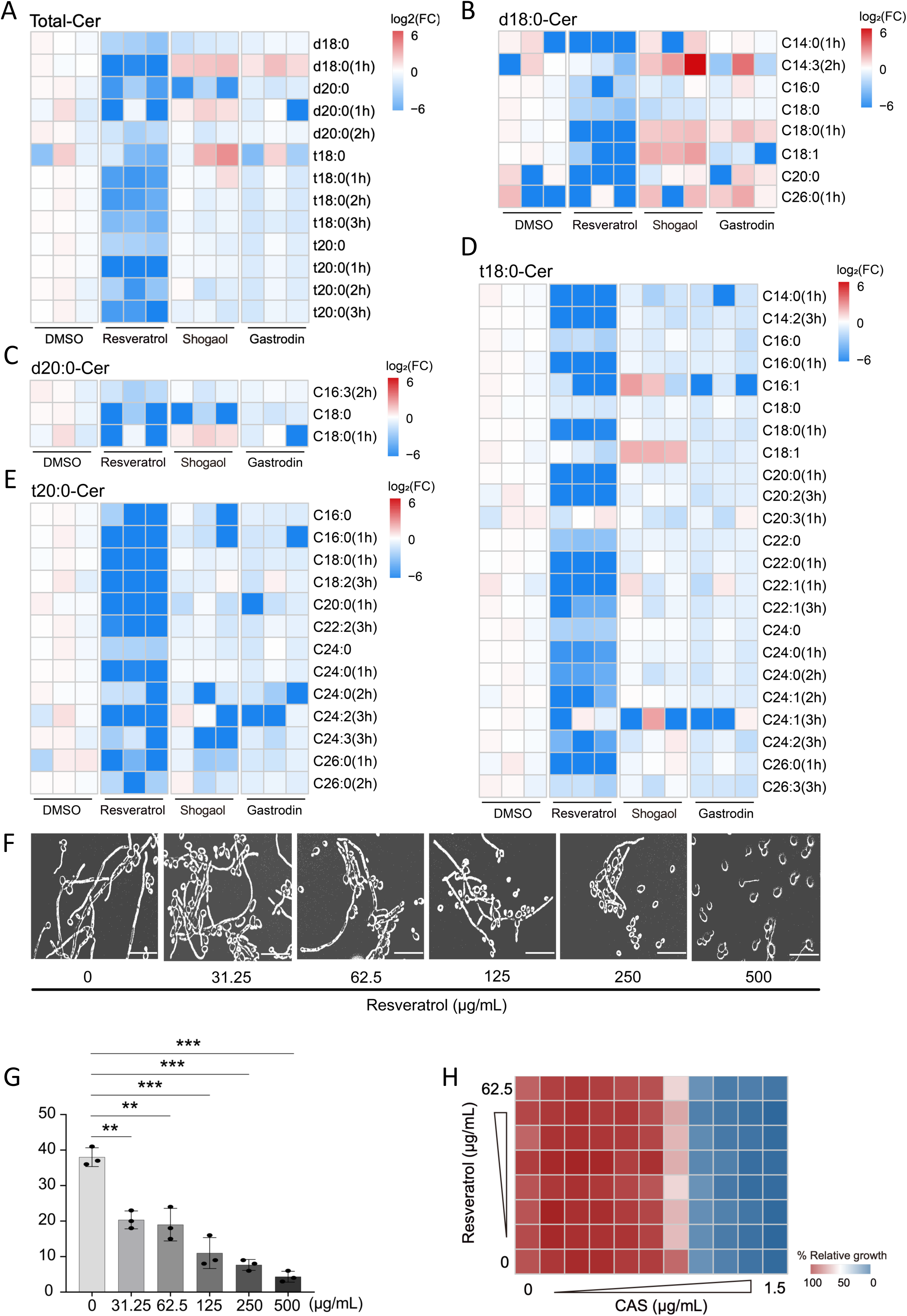
Resveratrol reduces hydroxy sphingolipid levels in *C. albicans* and exerts synergistic antifungal effects with caspofungin. (A) Heatmap showing the relative abundance of total ceramide species in *C. albicans* following 12 h treatment with DMSO, Resveratrol, Shogaol, or Gastrodin, each applied at 30 μg/mL. Ceramides are classified by long-chain base type, dihydrosphingosine (d-18:0 and d-20:0) or phytosphingosine (t-18:0 and t-20:0), and fatty acid hydroxylation status (1h/2h/3h, indicating the number of hydroxyl groups).Data are presented as log₂-transformed fold changes relative to the DMSO control (B–E) Heatmaps showing the abundance of species with different fatty acid chain lengths under the same treatment conditions as in (A). (B) d18:0 dihydroceramides. (C) d20:0 dihydroceramides. (D) t18:0 phytoceramides. (E) t20:0 phytoceramides. 1h, 2h, and 3h denote the number of hydroxyl groups on the fatty acid chain. (F-G) Hyphal formation of wild-type *C. albicans* was detected after culture in Spider medium supplemented with different concentrations of Resveratrol at 37°C for 24 h. The applied concentrations were 0, 31.25, 62.5, 125, 250 and 500 μg/mL. Representative microscopic images are shown in (F), scale bar = 10 μm. (G) shows the proportion of hyphal cells calculated by counting no fewer than 100 cells in each group. (H) Checkerboard assay was performed to assess the interaction effect between Resveratrol and caspofungin (CAS) against *C. albicans*. All data are presented as mean ± SD. Two-tailed unpaired *t*-tests were adopted for statistical analysis in (G). Statistical differences were defined as \**p* < 0.05, \*\**p* < 0.01 and \*\*\**p* < 0.001, with a sample size of n = 3.

Building on this metabolic disruption, we next examined the impact of Resveratrol on hypha formation. Resveratrol suppressed hyphal development in a dose-dependent manner (Figures 8F,G), a phenotype highly consistent with that of the *scs7*Δ/Δ mutant.

To test whether Resveratrol could potentiate *C. albicans* susceptibility to caspofungin by inhibiting Scs7, we assessed the combined antifungal activity of Resveratrol and caspofungin using a checkerboard assay. The combination of Resveratrol and caspofungin markedly enhanced the inhibitory effect against *C. albicans* compared with either agent alone (Figure 8H). Collectively, these findings indicate that Resveratrol inhibits Scs7 activity, reduces hydroxylated ceramide levels, impairs hypha formation, and sensitizes *C. albicans* to caspofungin. These findings provide experimental support for Resveratrol as a candidate for combination therapy targeting sphingolipid metabolism.

## Discussion

In this study, by integrating genetic screening with *in vitro* assays and an *in vivo* skin infection model, we identify the sphingolipid hydroxylase SCS7 as a central regulator of caspofungin resistance and cutaneous virulence in *Candida albicans*. Furthermore, combining virtual screening with experimental validation, we find Resveratrol as a natural scaffold targeting Scs7.

Prior studies established that modulating fungal sphingolipid synthesis or composition can significantly influence fungal drug resistance and pathogenicity [28,31–36]. For instance, Sharma et al. showed that deletion of the sphingolipid fatty acid elongase genes *FEN1* and *SUR4* in *Saccharomyces cerevisiae* specifically increased sensitivity to the polyene amphotericin B [37], a finding later confirmed for the orthologous genes *CaFEN1*/*CaFEN12* in *Candida albicans* [38]. Similarly, Hassoun et al. demonstrated in an intestinal colonization model that elevated phytosphingosine levels reduce caspofungin susceptibility[39]. Extending this paradigm, our data demonstrate that deletion of the sphingolipid α-hydroxylase gene *SCS7* markedly enhances *C. albicans* sensitivity to the echinocandin caspofungin, suggesting that α-hydroxylation of the fatty acid chain is a critical regulator of fungal drug resistance and host interaction.

The fungal cell wall is a critical bulwark against stress [19,40]. Previous work linked sphingolipid depletion to compromised cell wall integrity in *S. cerevisiae* [17]. We extend these findings by showing that *SCS7* deletion disrupts cell wall homeostasis in *C. albicans*. Mechanistically, sphingolipid depletion is known to perturb lipid raft architecture, blocking the proper localization of cell wall synthases like Gas1p [41]. Septins act as molecular scaffolds that, in a sphingolipid-dependent manner, couple lipid raft domains to the recruitment of downstream cell wall biosynthetic enzymes [42]. We propose that loss of hydroxylated sphingolipid levels upon *SCS7* deletion perturbs plasma membrane lipid raft organization, consequently interfering with the correct localization and function of cell wall biosynthetic proteins and ultimately leading to aberrant cell wall assembly.

Beyond its role in maintaining cell wall homeostasis, sphingolipid modification-mediated remodeling of lipid raft microdomains also participates in regulating the virulence-associated morphogenetic transition in *Candida albicans* [43,44]. The yeast-to-hypha transition is a key virulence trait that underpins host adhesion, tissue invasion, and biofilm formation [45], and the cAMP-PKA signaling pathway constitutes the central signaling axis governing this morphogenetic switch [46,47]. The adenylyl cyclase Cyr1, a critical catalytic enzyme of this pathway, is localized to the plasma membrane, and its biological activity may be dependent on lipid raft microdomains enriched in sphingolipids and sterols [48]. Previous studies have established that hydroxylation of the sphingoid base of phytoceramide is an important structural determinant for maintaining membrane order [49]. Our data show that *SCS7* deletion reduces intracellular hydroxylated sphingolipid levels and decreases cAMP production in *C. albicans*. We therefore propose that the loss of hydroxylated sphingolipids likely disrupts the structural integrity of plasma membrane lipid rafts, thereby interfering with the proper membrane localization and catalytic function of Cyr1. This, in turn, leads to reduced cAMP levels, impaired activation of the downstream PKA signaling cascade, and ultimately suppression of hyphal development and the virulence transition in *C. albicans*.

By homology to mammalian FA2H, which has been reported to act on free fatty acids, fungal Scs7 is thought to differ in that it modifies sphingolipid-linked acyl chains [50,51]. This divergence was not experimentally validated here but informed our targeting strategy. This functional divergence underscores the therapeutic potential of Scs7 as a selective antifungal target. Leveraging this, we identified resveratrol as a Scs7 inhibitor that synergizes with caspofungin. Sphingolipidomic profiling confirmed that Resveratrol significantly downregulates hydroxylated phytoceramides, further validating its on-target inhibitory effect on Scs7. However, the binding affinity between resveratrol and Scs7 has not yet been experimentally determined, and evidence for their direct interaction therefore remains to be established. Notably, unlike *SCS7* gene deletion, Resveratrol treatment did not lead to an accumulation of non-hydroxylated ceramides, suggesting that this compound may not exclusively target Scs7 but rather acts on multiple regulatory nodes within the sphingolipid metabolic network.

In summary, we establish sphingolipid hydroxylase as a critical determinant of caspofungin resistance and cutaneous infection by *C. albicans*. We validate Scs7 as a druggable target and demonstrate that Resveratrol inhibits its function to potentiate caspofungin efficacy. This work not only deepens our mechanistic understanding of how sphingolipid metabolism regulates fungal drug resistance, but also provides a theoretical and experimental foundation for developing novel antifungal strategies targeting *SCS7*. Our *in vivo* validation is currently limited to the murine cutaneous candidiasis model. Extending our findings to invasive systemic candidiasis and other clinically relevant infection settings will further enhance the translational potential of this target. Meanwhile, structural optimization of resveratrol to improve its Scs7 inhibitory efficacy will lay a tangible foundation for advancing antifungal drug development.

## Materials and Methods

### Microbial strains

All *Saccharomyces cerevisiae* strains used in this study were constructed by standard genetic techniques [52]. *Candida albicans* mutant strains were generated in the SN152 background using a fusion PCR-based homologous recombination strategy, in which the two alleles of the target gene were sequentially replaced with the auxotrophic markers *Cm.LEU2* and *Cd.HIS1* to yield homozygous deletion mutants. Complemented strains were constructed by a PCR-mediated in situ complementation strategy, in which the target gene open reading frame together with its upstream and downstream flanking regions was fused with the *Cd.ARG4* marker and introduced into the deletion mutant. The *C. albicans* wild-type strain SN250 served as the wild-type control. Detailed information on all strains is provided in Tables S1 and S2. *S. cerevisiae* and *C. albicans* strains were routinely cultured in YPD medium (1% yeast extract, 2% peptone, 2% glucose) at 30°C.

### Mice

All animal experiments were conducted in strict accordance with the Regulations for the Administration of Laboratory Animals issued by the Ministry of Science and Technology of the People’s Republic of China. The study protocol was reviewed and approved by the Institutional Animal Care and Use Committee (IACUC) of Nanchang University (Approval No. NCULAE-20251230001). Female SPF-grade BALB/c-nu nude mice (6–8 weeks old) were housed under pathogen-free conditions at 21°C with 50%–70% relative humidity and a 12-hour light/dark cycle, with free access to food and water. All experimental procedures were performed in accordance with IACUC approved protocols. No sex-based analyses were conducted in this study.

### Spot assay

Test strains were grown overnight in YPD medium at 30°C to logarithmic phase. Starting from an optical density of 1 OD600 unit/mL, ten-fold serial dilutions were prepared. Aliquots of the diluted cell suspensions were spotted onto drug-containing YPD agar plates and incubated at 30°C for two days, after which cell growth was assessed.

### Minimum inhibitory concentration assay

Antifungal susceptibility was evaluated using the broth microdilution method in accordance with the M27-A4 standard protocol of the Clinical and Laboratory Standards Institute (CLSI). Briefly, fungal cells were resuspended in RPMI 1640 medium, adjusted to the appropriate concentration, and added to 96-well microplates. Serially diluted antifungal agents were then added, and the plates were incubated at 35°C for 24–48 h before endpoint reading.

### Filamentation assay

*Candida albicans* cells grown to mid-log phase in fresh YPD medium at 30°C were adjusted to 2 × 10⁵ CFU/mL for hyphal induction. For liquid culture, cells were resuspended either in RPMI 1640 medium supplemented with 10% fetal bovine serum (FBS) or in Spider liquid medium and incubated at 37°C for 24 h. Hyphal morphologies were observed and imaged under an upright microscope, and hyphal morphotypes were subsequently quantified using ImageJ software. For solid culture, cells were inoculated onto RPMI 1640 or Spider agar plates containing 10% FBS and incubated at 37°C for 7 days, after which colony morphologies were photographed and recorded.

### Biofilm formation assay

Sterile 12-well plates were pre-treated with fetal bovine serum at 37°C overnight. *C. albicans* cells (1 × 10⁶ CFU) suspended in Spider medium were added to each well and incubated at 37°C for 90 min. Non-adherent cells were removed by washing with PBS, and fresh Spider medium was added. After further incubation at 37°C for 24 h, cells were washed again with PBS and fixed with methanol for 30 min. Methanol was discarded, and 2 mL of 1% crystal violet solution (Cat# C8470, Beijing Solarbio Science & Technology Co., Ltd.) (in 20% ethanol) was added to each well, followed by staining on an orbital shaker for 1 h. After washing until colorless, 2 mL of glacial acetic acid was added to each well, and the plates were shaken for 30 min to elute the stain. Finally, 200 μL of the destaining solution was transferred to a 96-well plate, and the absorbance was measured at 620 nm using a microplate reader.

### Cell wall chitin staining

*C. albicans* cells from overnight culture were collected by centrifugation, washed twice with PBS, and adjusted to a concentration of 1 × 10⁷ CFU/mL. One milliliter of the cell suspension was transferred to a sterile 1.5 mL microcentrifuge tube. Cells were pelleted, resuspended in 1 mL of 4% paraformaldehyde, and fixed for 1 h at room temperature. After fixation, cells were again pelleted, washed three times with PBS, and resuspended in 500 μL of 30 μg/mL Calcofluor White (CFW) solution (Cat# SL7204, Coolaber). The suspension was incubated at 37°C for 1 h in the dark with shaking. Following incubation, cells were centrifuged, the supernatant was removed, and the pellet was resuspended in 100 μL of PBS. A 10 μL aliquot was placed onto a clean glass slide, covered with a coverslip, and examined under a laser confocal microscope to record fluorescence.

### Intracellular cAMP assay

*C. albicans* cells were cultured in Spider medium at 37°C for 24 h, then collected and washed with PBS. Intracellular cAMP levels were measured using the cAMP-Glo™ Assay kit (Cat# V1501, Promega) according to the manufacturer’s instructions.

### RNA extraction and quantitative real-time PCR

Total RNA was extracted from *C. albicans* and mouse skin tissues using the RNAiso Plus reagent (Cat# 9109, Takara Bio). Real-time PCR and quantitative analysis were performed using SYBR Green Real-Time PCR Master Mix (Mei5 Biotechnology) and a qTOWER³ G Real-Time PCR System (Biometra). Transcript levels of target genes were normalized to 18S rRNA for *C. albicans* and to *Actb* for mouse samples. Primer sequences are provided in Table S3.

### Lipid extraction and ceramide analysis

Lipid extraction and ceramide analysis were performed following our previously described protocol with minor modifications [53,54]. Briefly, logarithmic-phase *C. albicans* cells were pelleted by centrifugation and subjected to lyophilization overnight. The dried material was then resuspended in 100 μL of water. A 10 μL aliquot was taken for protein quantification, while the remaining suspension was mixed with 1.5 mL of extraction solvent (methanol:chloroform = 2:1, v/v) and 10 μL of 2.5 μmol/L internal standard solution (LIPID MAPS™ Sphingolipid Mix II, Avanti Polar Lipids, Alabaster, AL, USA, #LM-6005). After vortexing, the mixture was incubated at 48°C overnight, then cooled to room temperature. Subsequently, 150 μL of 1 M KOH in methanol was added, and the mixture was incubated at 37°C for 2 h with shaking. After cooling, 6 μL of glacial acetic acid was added for neutralization. Then, 2 mL of water and 1 mL of chloroform were added, and the mixture was vortexed and centrifuged. The lower chloroform phase was collected, and the remaining solution was re-extracted with another 1 mL of chloroform. The combined chloroform phases were evaporated to dryness using a rapid vacuum concentrator. The residue was dissolved in 250 μL of a methanol–water–acetonitrile–isopropanol mixture (1:1:1:3, v/v/v/v containing 7 mM ammonium acetate) for LC-MS/MS analysis. Data were normalized to protein content and internal standard.

An ultra-performance liquid chromatography (UPLC) system (Shimadzu, Kyoto, Japan) coupled with a Triple Quad™ 5500+ QTRAP™ mass spectrometer (AB Sciex, Framingham, MA, USA) was used for sphingolipid analysis. The UPLC system was equipped with a binary pump, an autosampler, and a C18 column (3.5 μm, 100 × 2.1 mm, Phenomenex). Mass spectrometry was conducted in positive electrospray ionization mode with multiple reaction monitoring (MRM). The MRM ion pairs (m/z) are summarized in Table S4. Mobile phase A consisted of methanol:water:acetonitrile (7 mM ammonium acetate) = 1:1:1 (v/v/v), and mobile phase B was isopropanol (7 mM ammonium acetate). The gradient program was 0–1.0 min, 50% B; 1.0–8.0 min, 50% to 98% B; 8.0–11.0 min, 98% B; 11.0–11.1 min, 98% to 50% B; 11.1–12.0 min, 50% B. The flow rate was 0.3 mL/min, and the injection volume was 5 μL. Sphingolipid quantification was performed by integrating peak areas for analytes and internal standards using SCIEX OS software (Version 3.0.0.3339). Data were normalized to internal standards and protein content.

### Murine skin colonization model

For skin colonization, fungal cells were prepared as described above, washed three times with PBS, and adjusted to a concentration of 5 × 10⁹ CFU/mL. A sterile cotton swab was used to apply 1 × 10⁹ cells to the dorsal skin of each mouse. Treatments were applied every two days. On day 14 of the experiment, dorsal skin tissues were collected and weighed. Spontaneous scratching behaviors of the colonized mice were observed and recorded for 30 min. Skin tissues were fixed, embedded, and sectioned, followed by hematoxylin and eosin (H&E) staining and periodic acid–Schiff (PAS) staining for histomorphological analysis.

### Virtual screening and molecular dynamics simulation

The three-dimensional structure of the Scs7 protein was predicted using AlphaFold 3. The receptor structure was preprocessed with the Protein Preparation Workflow module in Maestro software (Schrödinger, New York, NY, USA), and potential binding sites were predicted using the SiteMap module. A library of molecules with reported antifungal activity was compiled from the literature, and their three-dimensional conformations were generated using the LigPrep module. High-throughput virtual screening (HTVS) and extra precision (XP) docking were performed sequentially using the Glide module, with threshold values set to −6.5 kcal/mol and −7.5 kcal/mol, respectively. The binding free energies of ligand–receptor complexes were subsequently calculated using the Prime-MMGBSA module. Molecules with binding free energies below −40 kcal/mol were selected, clustered based on two-dimensional structural similarity, and further filtered by removing those with unreasonable docking conformations, ultimately identifying candidate compounds. For the selected Scs7-small molecule complexes, a membrane environment was built using the PPM 3.0 server. The systems were then solvated, neutralized with chloride ions, and subjected to molecular dynamics simulations under the NPT ensemble using the Desmond module to evaluate the stability and dynamic behavior of the complexes.

### Checkerboard assay

For the checkerboard assay, Resveratrol (Cat# T1558, TargetMol), Shogaol (Cat# T2891, TargetMol), Gastrodin (Cat# T2S0223, TargetMol), or caspofungin (Cat# GC17230, GLPBIO) were dissolved in YPD liquid medium to prepare 400× stock solutions (with an equal volume of DMSO as control), giving a final concentration of 4×. These stock solutions were then added to designated wells of a 96-well plate so that the final concentration of each compound was 2×. Overnight cultures of the test strains were diluted with YPD medium to approximately 2 × 10⁴ CFU/mL, and the diluted suspension was added to each well to achieve the working concentrations of the drugs. After incubation at 30°C for 36 h, the OD600 was measured using a microplate reader, and the results were normalized to the DMSO control wells.

### Quantification and statistical analysis

All experiments in this study were performed at least three biological replicates. Quantitative data are presented as the mean ± standard deviation. Comparisons between two groups were performed using Student’s *t*-test, and comparisons among multiple groups were performed using one-way analysis of variance (ANOVA) followed by pairwise comparisons. GraphPad Prism (Version 10.4.0) and R (Version 4.5.3) were used to generate data plots and perform statistical tests. Significance levels are indicated as \**p* < 0.05, \*\**p* < 0.01, and \*\*\**p* < 0.001.

## Resource availability

### Lead contact

Requests for further information and resources should be directed to and will be fulfilled by the lead contact, Jingjing Duan.

### Materials availability

This study did not generate any new, unique reagents. Requests for microbial strains, including the mutant strains generated in this study, should be addressed to the lead contact and are subject to a material transfer agreement.

### Data availability

All data needed to evaluate the conclusions of this study are present in the paper or the supplemental information. The public transcriptomic dataset re-analyzed in this study is available in the NCBI Gene Expression Omnibus (GEO) repository under accession number GSE327269. No new datasets were generated in the present work. Any additional information required to reanalyze the data reported in this paper is available from the lead contact upon request.

## Acknowledgments

We thank Dr. Zhenying Hu (Nanchang Medical College) for his technical expertise and suggestions regarding the sphingolipidomic profiling of *Candida albicans*, and Dr. Yuping Zhang (Shanghai Institute of Immunity and Infection, Chinese Academy of Sciences) for his guidance on *C. albicans* culture and related experiments. This work was supported by the National Natural Science Foundation of China (32471359) and the Natural Science Foundation of Jiangxi Province (20232ACB205002 and 20242BAB23056) to Jingjing Duan.

## Author contributions

The study was conceived by J.D., X.C., and S.M. *In vitro* experiments and mouse skin infection experiments were performed by X.C. and S.M., and J.P. Sphingolipid extraction from *Candida albicans* was performed by X.C. and S.M. *Saccharomyces cerevisiae* and *Candida albicans* mutant strains were constructed by X.C. and Z.D. Q.Z., L.Z., and X.H. provided *C. albicans* strains and offered suggestions on related experiments. Virtual screening of small molecules was performed by B.Y. Mouse skin tissue collection was performed by X.C., S.M., and M.Z. Data interpretation, analysis, and visualization were performed by J.D., X.C., and S.M. The manuscript was written by S.M. and J.D. All authors have read and approved the manuscript.

---

**Figure S1.**
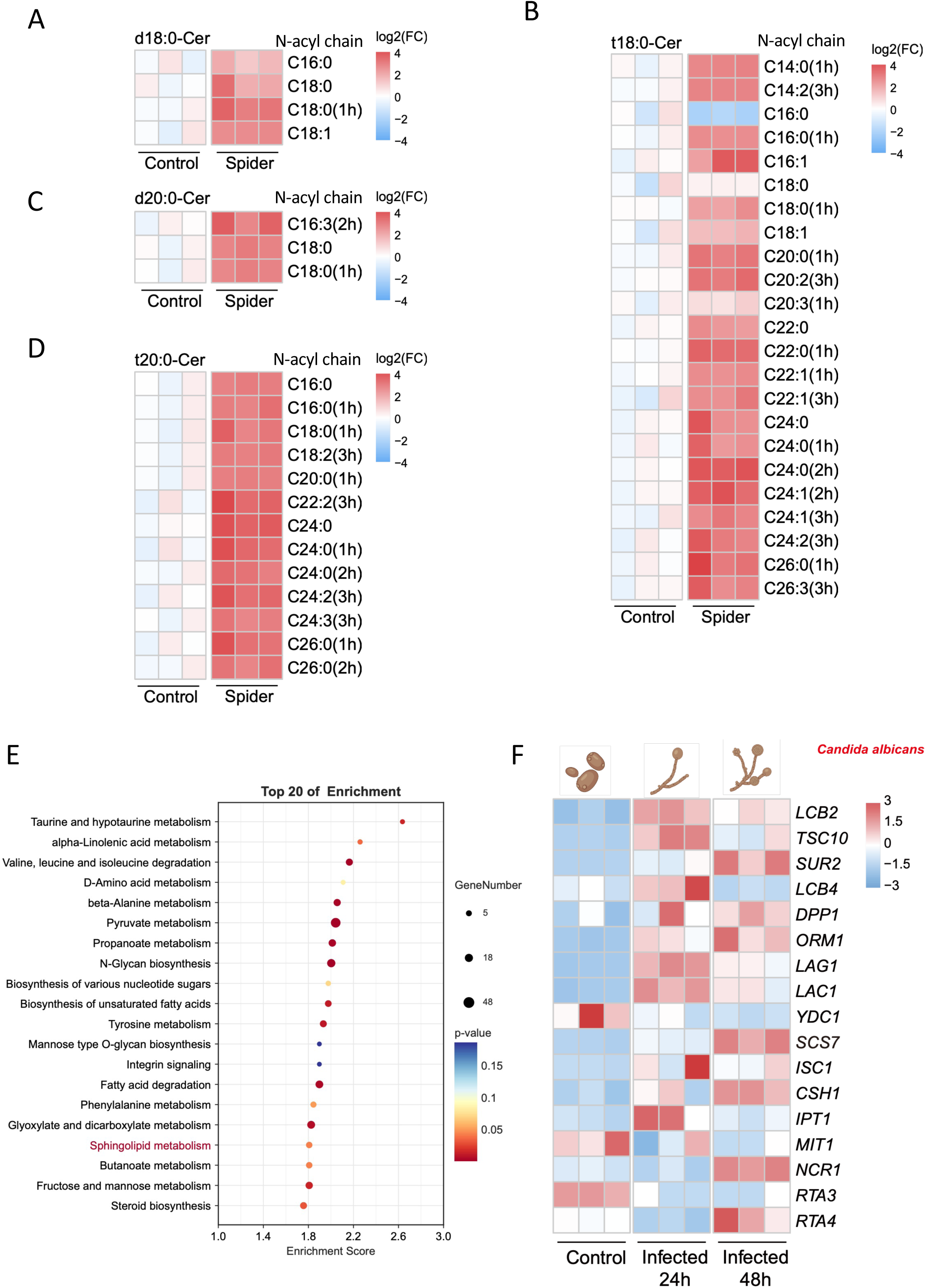
Sphingolipidomic profiling and transcriptomic analysis of *C. albicans* during hyphal induction and in vivo infection. (A–D) Heatmaps showing the log₂ fold change (FC) of ceramide species in *C. albicans* yeast-form (Control) and hyphae cells (Spider). Ceramides are classified by long-chain base type, dihydrosphingosine (d18:0 and d20:0) or phytosphingosine (t18:0 and t20:0), and fatty acid hydroxylation status (1h/2h/3h, indicating the number of hydroxyl groups). (A) d18:0 dihydroceramide. (B) d20:0 dihydroceramide. (C) t18:0 phytoceramide. (D) t20:0 phytoceramide. Data are presented as log_2_ fold change (FC) relative to the Control. (E) KEGG pathway enrichment analysis showing the top 20 enriched metabolic pathways in *C. albicans* at 48 h post-infection in a mouse kidney infection model, based on the public transcriptomic dataset GSE327269. (F) Heatmap showing the expression of sphingolipid metabolism-related genes in *C. albicans* isolated from infected mouse kidneys at 0 h (uninfected control), 24 h, and 48 h post-infection, derived from the same dataset above. Color intensity represents Z-score normalized expression levels (blue, downregulated; red, upregulated).

**Figure S2.**
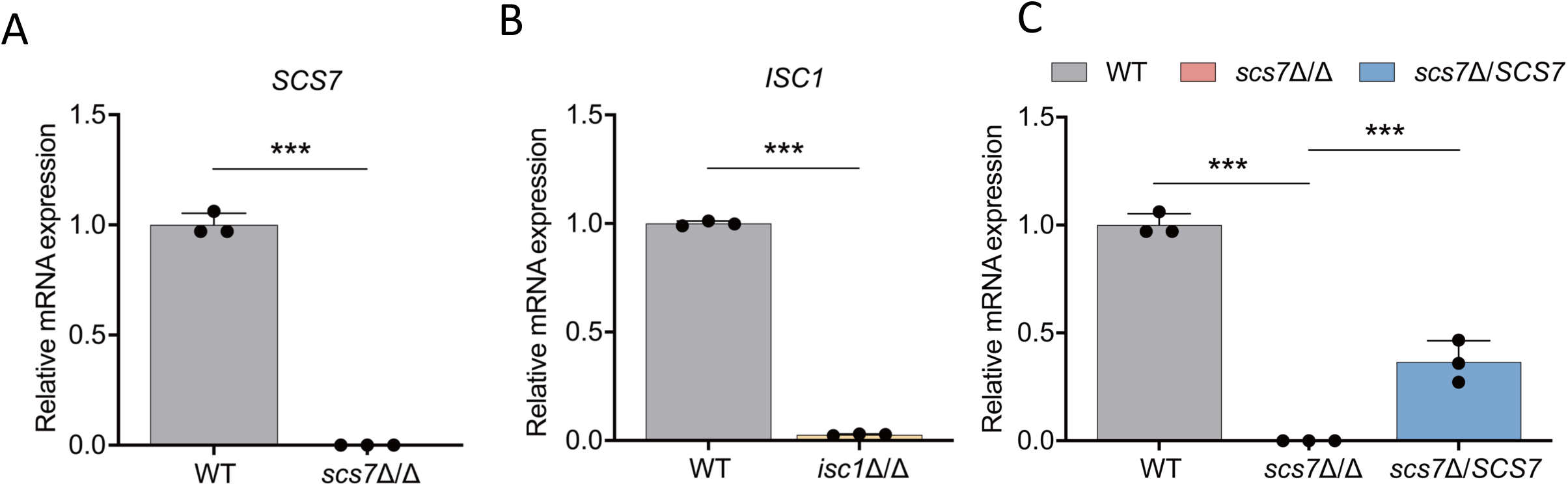
**Transcriptional confirmation of *SCS7* and *ISC1* deletion in *C. albicans*** (A-C) Relative mRNA expression levels of target genes in *scs7*Δ/Δ, *isc1*Δ/Δ, and *scs7*Δ/*SCS7* strains. Data are presented as mean ± SD and were analyzed by two-tailed unpaired *t*-tests (A–C), with significance levels indicated as \**p* < 0.05 and \*\**p* < 0.01 (n = 3).

**Figure S3.**
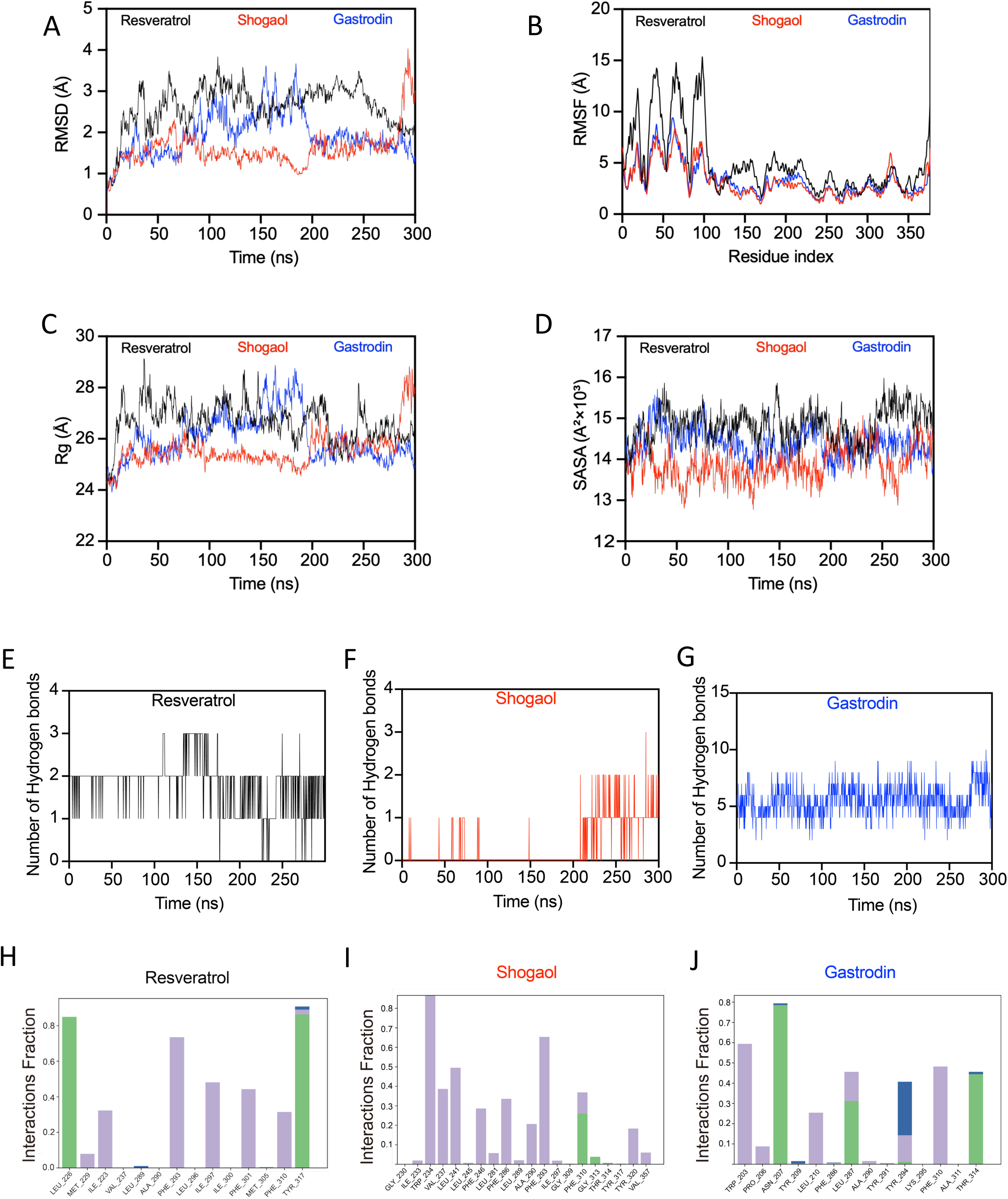
Molecular dynamics simulation analyses of Scs7–compound complexes: Scs7–Resveratrol (black), Scs7–Shogaol (red), and Scs7–Gastrodin (blue). (A–D) Key structural stability metrics: (A) RMSD of Cα atoms; (B) RMSF of residues; (C) Radius of gyration (Rg); (D) Solvent–accessible surface area (SASA). (E–G) Number of protein–ligand hydrogen bonds formed during the 300 ns simulation: (E) Scs7–Resveratrol, (F) Scs7–Shogaol, (G) Scs7–Gastrodin. (H–J) Temporal evolution of non–covalent interactions (hydrogen bonds (green), water bridges (blue), hydrophobic contacts (purple)) between Scs7 and (H) Resveratrol, (I) Shogaol, or (J) Gastrodin.

